# Reducing mitochondrial reads in ATAC-seq using CRISPR/Cas9

**DOI:** 10.1101/087890

**Authors:** Lindsey Montefiori, Liana Hernandez, Zijie Zhang, Yoav Gilad, Carole Ober, Gregory Crawford, Marcelo Nobrega, Noboru Jo Sakabe

## Abstract

ATAC-seq is a high-throughput sequencing technique that aims at identifying DNA sequences located in open chromatin. Depending on the cell type, ATAC-seq may yield a high number of mitochondrial sequencing reads (~20-80% of the reads). As the regions of open chromatin of interest are usually located in the nuclear genome, mitochondrial reads are typically discarded from the analysis. To decrease wasted sequencing, we performed targeted cleavage of mitochondrial DNA using CRISPR/Cas9 and 100 mtDNA-specific guide RNAs. We also tested a modified ATAC-seq protocol that does not include detergent in the cell lysis buffer. Both treatments resulted in considerable reduction of mitochondrial reads (1.7 and 3-fold, respectively). The removal of detergent, however, resulted in increased background and fewer peaks identified. The highest number of peaks and highest quality data was obtained by preparing samples with the original ATAC-seq protocol (using detergent) and treating them with anti-mitochondrial guide RNAs and Cas9. This strategy could lead to considerable cost reduction and improved peak calling when performing ATAC-seq on a moderate to large number of samples and in cell types that contain a large amount of mitochondria.

## Introduction

ATAC-seq aims at identifying DNA sequences located in open chromatin, i.e., genomic regions whose chromatin in not densely packaged and that can be more easily accessed by proteins than closed chromatin. The ATAC-seq technique makes use of the Tn5 transposase, an optimized hyperactive transposase that fragments and tags the genome with sequencing adapters in regions of open chromatin(1). The output of the experiment is millions of DNA fragments that can be sequenced and mapped to the genome of origin for identification of regions where sequencing reads concentrate and form “peaks”.

While ATAC-seq often generates high-quality data with low background, certain cell types and tissues yield an enormous fraction (typically 20-80%) of unusable sequences of mitochondrial origin. In order to reduce the amount of wasted sequencing reads, targeted cleavage of DNA fragments has recently been used to deplete mitochondrial ribosomal RNA-derived fragments in RNA-sequencing libraries(2). In another study, Wu *et al*. targeted the mitochondrial genome in ATAC-seq experiments using 114 guide RNAs (gRNAs) and observed a ~50% decrease in mitochondrial reads and no adverse modification of the readenrichment pattern(3).

To test the efficacy of this approach, we designed 100 gRNAs targeting the human mitochondrial chromosome every ~250 base pairs (bp) and treated ATAC-seq sequencing libraries with these gRNAs and Cas9 enzyme(4), hereafter referred to as anti-mt CRISPR. We compared this method to a modified ATAC-seq protocol that also aims at reducing mitochondrial reads by removing detergent from the cell lysis step, which is believed to prevent lysis of the mitochondrial membrane (Supplemental File 1).

We observed that while both methods considerably reduced the number of mitochondrial reads sequenced, each method displayed different effects on the number of peaks called. Whereas the removal of detergent from the lysis buffer had the largest effect in reducing mitochondrial reads, it resulted in decreased quality of the ATAC-seq libraries, as measured by the number of peaks called at a given sequencing depth, the total number of reads in peaks, and the fraction of transcription start sites (TSS’s) and enhancers identified. Conversely, in addition to decreasing the number of mitochondrial reads, the anti-mt CRISPR treatment also resulted in a greater number of peaks, a greater number of reads in peaks, and higher overlap of peaks with TSS’s and enhancers.

Performing anti-mt CRISPR requires the one-time purchase of gRNA template oligos, as well as purchase of the Cas9 enzyme. The gRNAs can be generated from template DNA oligos indefinitely and shared as a community resource, potentially trivializing the up-front cost. Cas9 enzyme as currently priced adds an additional US $20 per ATAC-seqsample. Thus, although there is an initial cost to treating ATAC-seq libraries, the reduction in wasted sequencing and potential for sharing of resources should bring the per-library cost down substantially when performing multiple ATAC-seq experiments or using cell types that yield a high fraction of mitochondrial reads.

## Material and methods

### Human lymphoblastoid cell line growth and harvesting

Human lymphoblastoid cell line NA19193 was obtained from Coriell Cell Repository. Cells were grown in RPMI 1640 medium lacking L-Glutamine (Corning), supplemented with 15% fetal bovine serum, 1% GlutaMAX (ThermoFisher) and 1% penicillin-streptomycin solution (ThermoFisher) at a density of 0.5×10^6^ to 1.0×10^6^ cells/mL. Cells were passaged every 2-3 days to maintain this density. Cells were harvested for ATAC-seq by centrifugation at 500 × g for 5 minutes at 4°C and resuspended in PBS. Cells were counted using a hemocytometer and 50,000 cells were immediately placed into a 1.5 mL Eppendorf tube for ATAC-seq.

### Preparation of ATAC-seq libraries

ATAC-seq libraries were generated according to the protocol of Buenrostro et al.(5) with minor changes. Instead of NEBNext High-Fidelity 2X PCR Master Mix, we used Q5 Hot Start High-Fidelity 2X Master Mix (New England Biolabs). Following PCR-amplification, instead of using a column to clean the reaction, we used a 0.8X Ampure bead purification and eluted the library with 20 μl nuclease-free water. For the ND samples, Igepal-CA630 was removed from the lysis buffer and replaced with water. One microliter of the library was used to run a high sensitivity Bioanalyzer to determine fragment size distribution and concentration.

### Anti-mitochondrial CRISPR/Cas9 treatment

To deplete ATAC-seq libraries of DNA fragments derived from the human (hg38) mitochondrial genome, 100 high-quality guide RNAs that specifically targeted the mitochondrial genome roughly every 250 base pairs were chosen using the gRNA design tool at http://crispr.mit.edu (full list of guide sequences is in Supplemental File 2). Full-length guide RNAs were designed according to Gu *et al*. (2) and generated from single-stranded oligo templates (Integrated DNA Technologies) according to Lin *et al*. (11). Briefly, each oligo consisted of the sequence 5’-TAATACGACTCACTATAG(N_20_)GTTTAAGAGCTATGCTGGAAACAGCATAGCAAGTTTAAATAAGGCTAGTCCGTTATCAACTTGAAAAAGTGGCACCGAGTCGGTGCTTTTTTT-3’ where N_20_ corresponds to the 20 nucleotide guide RNA seed sequence. The PAM sequence would occur at the 3’ end of the N_20_ sequence. Oligos were purchased as a 200 picomole plate from Integrated DNA Technologies and received as a lyophilized pool. They were resuspended in 1 mL TE1.0 buffer (10 mM Tris-HCl, 0.1 mM EDTA). A Nanodrop was used to determine the concentration of the oligos and 8 ng were used as template for PCR to make them double-stranded. The PCR reaction consisted of 4 μl 5X HF Buffer (New England Biolabs), 0.4 μl 10 mM dNTPs, 1 μl of each 10 μM primer (For: 5’-TAATACGACTCACTATAG, Rev: 5’-AAAAAAAGCACCGACTCGGTGC), 0.2 μl Phusion High-Fidelity DNA Polymerase (New England Biolabs) and nuclease-free water to a final volume of 20 μl. Thermocycler conditions were 98°C for 30 s, followed by 30 cycles of 98°C for 10 s, 56°C for 10 s, 72°C for 10 s, and then a final extension of 72°C for 5 minutes. The reaction was cleaned using a Qiagen MinElute Purification kit and eluted in 10 μl of nuclease-free water. Enough PCR reactions were performed to obtain 1 μg of double-stranded template (should be in a volume of less than 8 μl). Transcription was carried out on 1 μg of template using the MEGAshortscript T7 Transcription kit (ThermoFisher) following manufacturer’s instructions and then cleaned with the MEGAclear Transcription Clean-Up kit (ThermoFisher). gRNAs were eluted from the column with 50 μl of RNase-free water and the concentration was determined using a Nanodrop, aliquoted and stored at −80°C.

We estimated that half of the DNA in each library was of mitochondrial origin, thus a 20 nM ATAC-seq library contained a mtDNA target concentration of 10 nM. Based on this value, 40, 100, 200, or 400 molar excess of Cas9 enzyme was used (New England Biolabs #M0386M) along with 100 molar excess gRNAs in a 30 μl reaction. The reaction was set up according to the protocol for Cas9 from *S. pyogenes* (New England Biolabs #M0386M). Briefly, the appropriate amounts of Cas9 enzyme and gRNAs were mixed with 3 μl of 10X Cas9 Buffer and water to a final volume of 22 μl. This was incubated at 25°C for 10 minutes and then 8 μl of the ATAC-seq library was added and the reaction was incubated at 37°C for one hour. For the two-hour treatment, the incubation was extended an additional hour; for the “Cas9 boost” treatment, the same amount of Cas9 enzyme was added after 1 hour of incubation and left for an additional hour. Reactions were subsequently treated with 1 μl of 20 mg/mL proteinase K for 15 minutes and purified using a Qiagen MinElute kit followed by elution in 10 μl nuclease-free water. Treated libraries were run on a high sensitivity Bioanalyzer chip to assess fragment size distribution and concentration (Supplemental Fig. S4). Because the multiplexing barcodes are added before treatment, for each batch of experiments, samples were sequenced on two lanes of an Illumina Hi-Seq 4000 instrument, separating anti-mt-CRISPR untreated and treated samples.

### Peak calling

Illumina reads were trimmed using cutadapt (12) and aligned to hg38 with Bowtie 2 version 2.2.3 (13) with default parameters. Reads with mapping quality lower than 10 were discarded. Mitochondrial reads and reads aligned to the same coordinates were removed. HOMER version 4.8.3 was run with 3 sets of parameters: (i) “default”: -style dnase -gsize 2.5e9, (ii) “ENCODE”: -localSize 50000—size 150—minDist 50 -fragLength 0 (https://www.encodeproject.org/pipelines/ENCPL035XIO/), (iii) “custom”:-gsize 2.5e9 -F 2 -L 2 -fdr 0.005 -region. MACS2 version 2.1.0 was run with 2 sets of parameters: (i) “default”:—nomodel—shift-100—extsize 200-q 0.01, (ii) “custom”:—nomodel-llocal 20000—shift-100—extsize 200.

### Fraction of TSS and enhancers intersecting peaks

Transcription start sites were obtained from the Gencode (8) GRCh38 basic set (ftp://ftp.sanger.ac.uk/pub/gencode/Gencode_human/release_24/gencode.v24.basic.annotation.gtf.gz), totaling 106,926 2 kb intervals centered on the TSS, and intersected with ATAC-seq peaks using bedtools intersect with the -u option (14). Epigenome Roadmap (9) 15-state ChromHMM coordinates were obtained from http://egg2.wustl.edu/roadmap/data/byFileType/chromhmmSegmentations/ChmmModels/coreMarks/jointModel/final/all.mnemonics.bedFiles.tgz. Coordinates were converted to hg38 using the UCSC Genome Browser liftOver tool and active enhancer (Enh7) states were intersected with peaks using bedtools with the -u option.

### Statistical tests

Due to the small number of replicates, we chose the more conservative one-sided unpaired Wilcoxon rank sum test to compare treatments in the boxplots shown (R statistical package version 3.3.1) (15). Student t-tests were in agreement with the results presented, yielding smaller P-values. Fold-differences of DT samples were calculated pairwise and the median was reported. Fold-differences of DT versus ND samples were calculated on the median of each group.

## Results

Our anti-mt CRISPR treatment consisted of 100 guide RNAs (gRNAs) targeting the human mitochondrial genome at regular intervals, which is usually densely covered by ATAC-seq reads generated from lymphoblastoid cell lines (LCLs), as shown in Fig. 1. The rationale was to cleave targeted DNA fragments in the sequencing library, rendering them unable to bind and amplify on the Illumina HiSeq flow cell. Similarly to Gu *et al*.(2) and Wu *et al*.(3), we chose to treat the final (PCR amplified) sequencing library with the gRNA/Cas9 mix instead of the unamplified tagmented DNA because of the small amount of DNA present in the sample at this earlier step. Although treating the samplesbefore PCR amplification might result in lower fractions of mitochondria, we chose the conservative approach of treating larger amounts of DNA to reduce technical variability.

**Figure 1.**
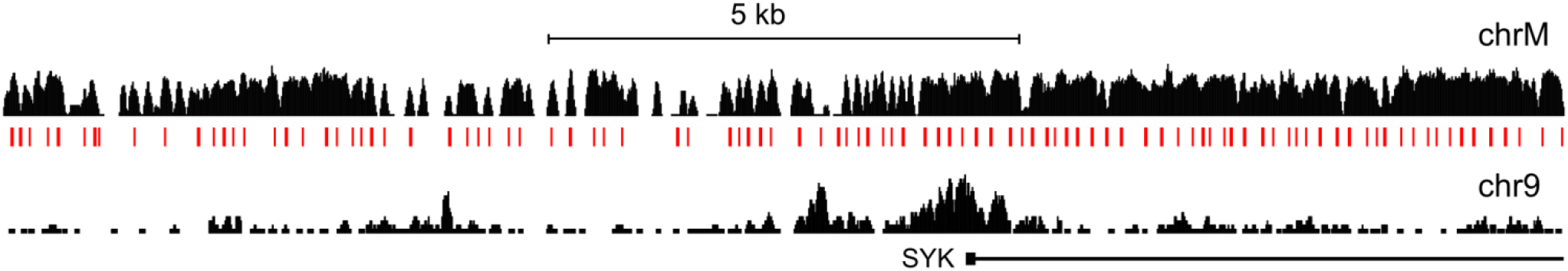
ATAC-seq read densities in the mitochondrial chromosome and one nuclear genome region. Top: The mitochondrial chromosome (chrM) is densely covered by uniquely mapped reads. Genomic location of the 100 mitochondrial guide RNAs (red tick marks) designed to target the human mitochondrial chromosome (top). Bottom: compare chrM to a 16.5 kb region of the nuclear genome (hg38, chr9:90,791,567-90,808,137). The chrM and chr9 tracks are shown in different height scales for easier visualization. Seven samples were pooled and 227 M reads were sampled.

To develop our anti-mt CRISPR treatment, we used 50,000 human LCLs per sample and generated a total of 27 pairs of ATAC-seq libraries for Illumina high-throughput sequencing according to the protocol of Buenrostro *et al*.(5)(Array Express accession number E-MTAB-5205 and Supplemental File 2). We split each of the 27 libraries into two equal parts, leaving one half untreated and treating the other half with 100 mitochondrial gRNAs and Cas9. Due to the single turn-over nature of Cas9, Gu *et al*.(2) used an excess of enzyme and of gRNA to deplete mitochondrial ribosomal DNA. Based on this notion, we used 100X Cas9 and 100X gRNA excess. We assumed 50% mtDNA fragments in the PCR-amplified ATAC-seq library to calculate exact amounts to be used in the treatment (See Methods).

We obtained between 9.8 M and 108.6 M reads per sample in four batches of experiments. Because different numbers of reads were sequenced from each sample due to imprecisions in DNA quantification and the number of multiplexed samples, we randomly sampled a fixed number of sequenced reads from each library in order to compare across samples. This approach allowed us to assess which library preparation method yielded the best results regardless of how it affected the number of aligned or usable (unique, non-mitochondrial) reads.

After aligning reads to the human genome, we removed mitochondrial reads and reads aligned to identical coordinates and called peaks using HOMER(6) and MACS2(7). Qualitatively similar results were obtained with both peak callers at three read depths (9.8 M (54 samples), 17 M (52 samples) and 21.9 M (47 samples)) and using different parameters to call peaks (Supplemental File 1). The results reported in the figures were obtained with MACS2 using custom parameters and 21.9 M sequenced reads. Results for all other read depths and parameters are presented in Supplemental Figs. S1 and S2 and Supplemental Tables S1 and S2 in Supplemental File 1.

Figure 2 shows the comparison between 14 ATAC-seq samples before and after treatment with anti-mt CRISPR, using the original ATAC-seq protocol that includes detergent (DT). Visual inspection of the data showed that the untreated and treated samples were similar (Fig. 2a), indicating that the treatment did not damage the samples. As expected, the anti-mt CRISPR treatment resulted in depletion of mitochondrial reads, while the number of reads in the nuclear genome increased (Fig. 2b).

**Figure 2.**
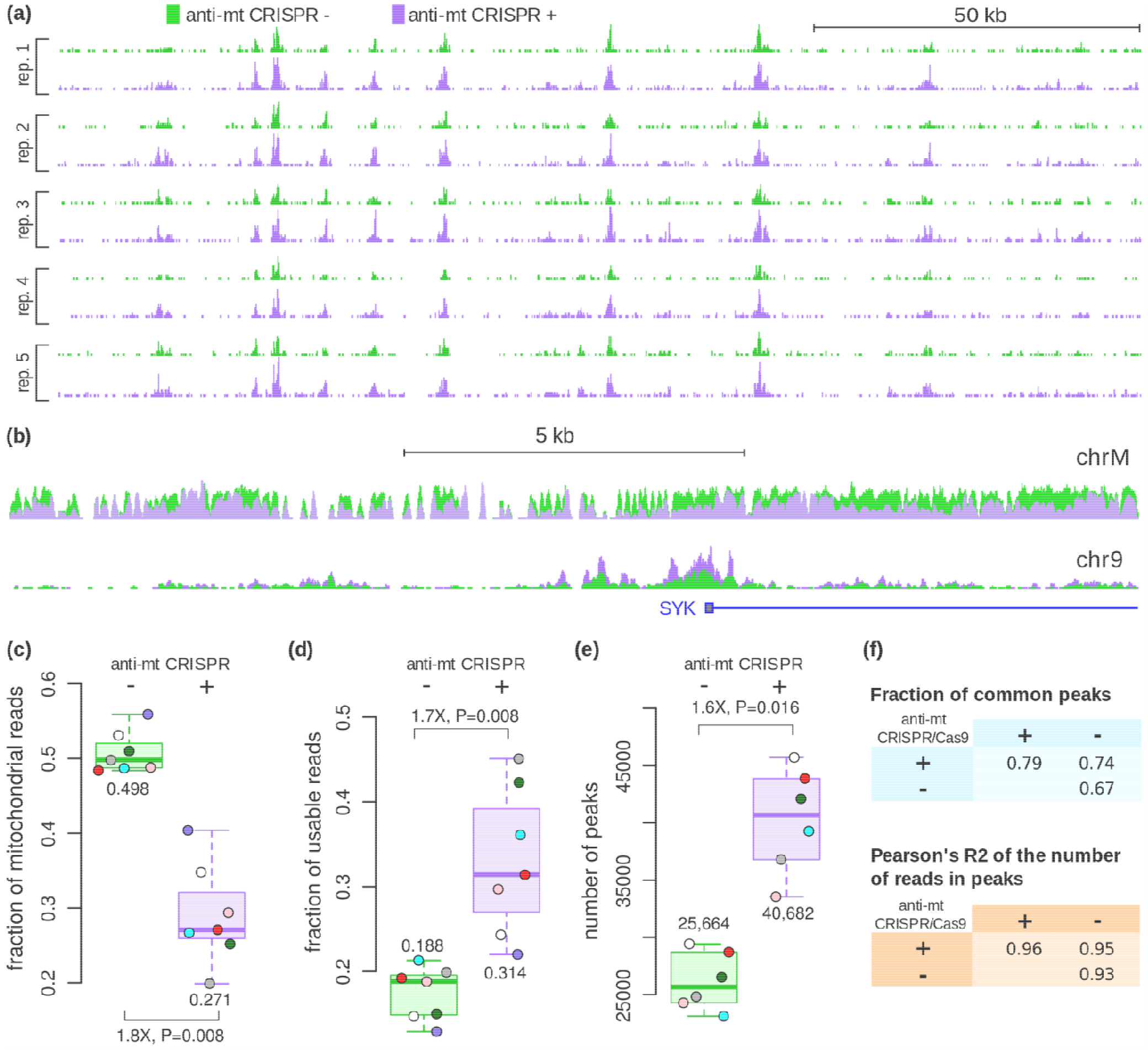
ATAC-seq was performed on human lymphoblastoid cells and half of each sample was left untreated (green) and the other half was treated with anti-mt CRISPR (purple). (a) Representative genomic region (hg38, chr2:74,425,417-74,586,546) showing read counts (usable reads) in 5 replicate pairs (DT) at the same sequencing depth of 21.9 M reads. (b) ATAC-seq reads in the mitochondrial chromosome and in a 16.5 kb region of chromosome 9 around the SYF promoter (same as Fig. 1). For each condition, all samples were pooled together and 227 M reads were sampled. (c) Treated samples yielded 1.8-fold fewer mitochondrial reads compared to untreated samples. (d) Accordingly, the number of unique, non-mitochondrial (usable) reads was 1.7-fold higher in treated samples than in their untreated counterparts. (e) At the same sequencing depth, 1.6-fold more peaks were called in the treated samples. Only 6 data points are shown because the treated halves of samples 18 and 19 (same batch) had only 14.5 M and 9.8 M reads each and were combined for improved peak calling. (f) Anti-mt CRISPR-treated samples shared a similar number of peaks with treated replicates and untreated samples. The top 20,000 peaks of each sample were used in this analysis. Comparison of peaks at the read count level also supports that peaks from treated samples do not substantially differ from untreated samples. Fold-differences were calculated on the medians.

At the same sequencing depth, the anti-mt CRISPR-treated samples yielded considerably less mitochondrial reads (Fig. 2c). This result is similar to the level of reduction of ~50% reported by Wu *et al*.(3). Consequently, more usable reads (non-mitochondrial reads with unique coordinates), were generated (Fig. 2d and Supplementary Fig. S1). The increased number of usable reads resulted in 50% more peaks in the treated halves of all samples (Fig. 2e and Supplementary Fig. S2), demonstrating the importance of removing excess mitochondrial reads from ATAC-seq samples.

One concern when treating samples with CRISPR/Cas9 was whether off-target gRNA/Cas9 activity would affect the data to a significant extent. To address this issue, we compared the percentage of peaks common across replicates (1 bp overlap) and across anti-mt CRISPR-treated and untreated samples. Figure 2f shows that the degree of overlap between untreated and treated samples was not smaller than the degree of overlap between replicates of the same condition, indicating that the anti-mt CRISPR treatment did not cause loss of peaks or create artefactual peaks. This observation is in accordance with a previous report that CRISPR treatment of sequencing libraries did not modify the read enrichment pattern(3). We also found evidence that the anti-mt CRISPR-treated samples identified more transcription start sites and enhancers than untreated samples (see below), indicating that mtDNA cleavage did not negatively affect the data.

### Effect of removing detergent from the original ATAC-seq protocol

Another method that has been used to reduce the fraction of mitochondrial reads is the complete removal of detergent from the cell lysis step of the ATAC-seq protocol (Supplemental File 1), which likely prevents disruption of the mitochondrial membrane and exposure of mtDNA to Tn5. We generated seven ATAC-seq samples with no detergent (ND) and observed several differences compared to the original protocol with detergent (DT).

Interestingly, the fraction of unique reads was considerably higher in ND samples compared to DT samples (56.5% vs 32.5%, respectively), which could reflect a lower fraction of mitochondrial fragments before PCR amplification of the sequencing library. In addition, the fraction of reads uniquely aligned to the genome was slightly higher in ND samples, compared to samples prepared with the original protocol (83.6% vs 74.6%, respectively). This difference is due to discarding reads that map to both mitochondrial and nuclear genomes (6% of ND reads and 17% of DT reads) in order to retain only uniquely aligned reads. Because we started our analyses with the same number of sequenced reads, these differences in mappability were accounted for in our comparisons.

Figure 3 shows that the removal of detergent had a pronounced depletive effect on mitochondrial reads compared to untreated DT libraries (Fig. 3a) and consequently increased the fraction of usable reads (Fig. 3b). Despite this 2.4-fold increase of ND usable sequences (0.45/0.19), the number of peaks called was higher by only 1.04-fold compared to untreated DT samples (Fig. 3c, 26,651/25,664). The mean fold-difference using other parameters to call peaks and read depths was higher at 1.2-fold, but still substantially lower than the increase in usable reads (Supplemental Table S1). This difference could be due to the increased background in ND samples (Figs. 3d and 3e), as suggested previously (Supplemental File 1). We considered background reads as reads that were not mitochondrial and were not in any ATAC-seq peak identified in any DT or ND sample. The lower signal/noise ratio in ND samples (Fig. 3d) provides an explanation for why fewer peaks wereidentified in ND samples. Thus, although removing detergent from the lysis buffer increased the overall number of non-mitochondrial reads, the background read coverage also increased, resulting in fewer peaks called at the same sequencing depth compared to the original protocol.

**Figure 3.**
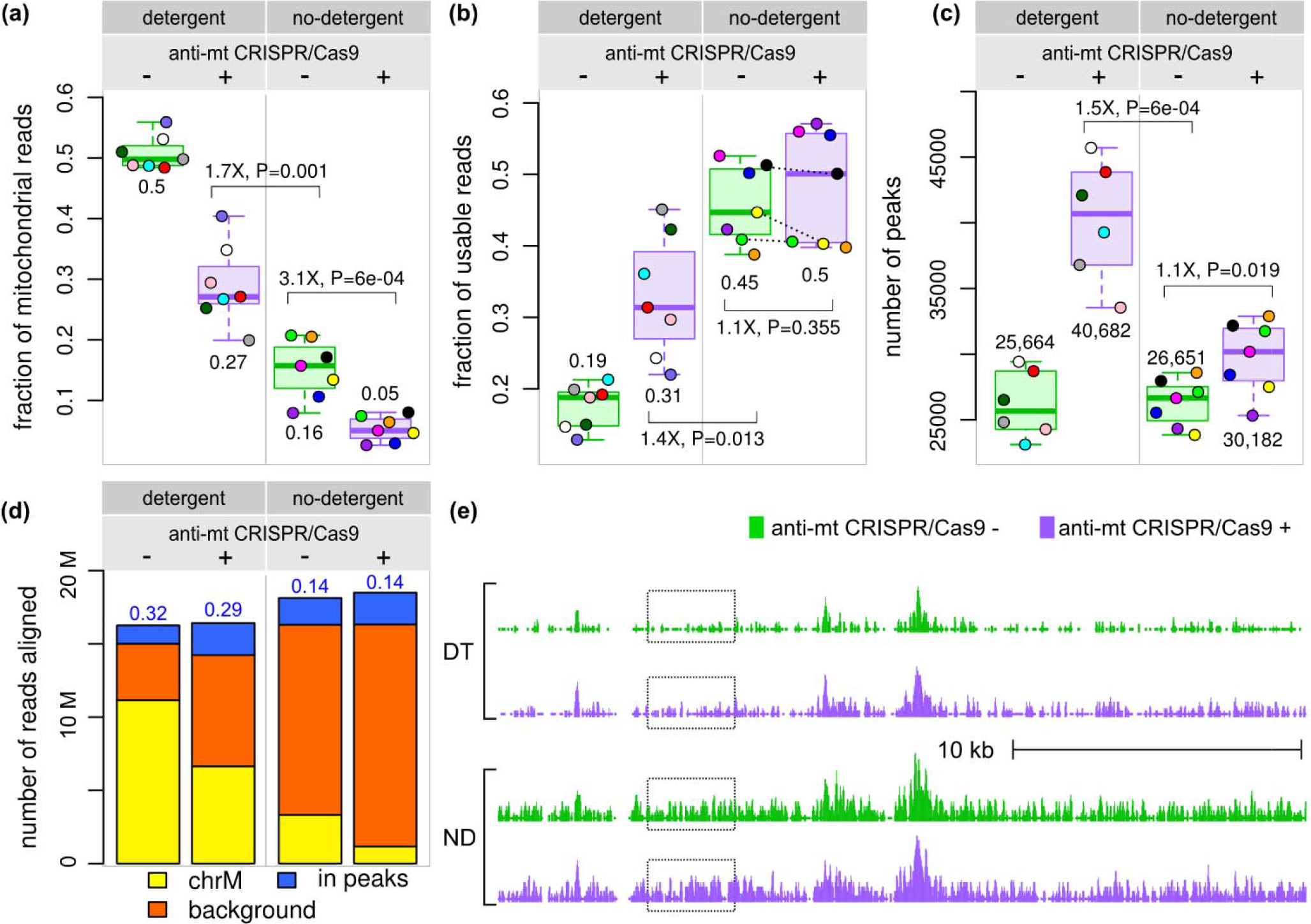
Effect of detergent removal from the ATAC-seq protocol. (a) The fraction of mitochondrial reads in samples prepared without detergent was considerably smaller than those prepared with the original protocol. Treatment with anti-mt DNA CRISPR led to further decrease of mitochondrial reads (3.1-fold). (b) The fraction of unique, non-mitochondrial reads was considerably higher when detergent was not used. Surprisingly, the anti-mt DNA CRISPR treatment had only marginal effect on the fraction of usable reads (1.1-fold increase). (c) At the same sequencing depth, only 1.1-fold more peaks were called in the ND treated samples. DT samples 18 and 19 were combined as in Fig. 2c. (d) ND samples displayed higher background (the number of non-mitochondrial reads outside peaks identified in any DT or ND sample). Numbers in blue above the bars are the ratio between number of reads in peaks and the number of background reads (signal/noise) (e) An example illustrating the higher background in ND samples, highlighted by the dashed boxes (chr6:420,146-448,555). Fold-differences calculated on medians.

Treating the ND samples with anti-mt CRISPR, i.e. combining anti-mt CRISPR treatment with the detergent-free lysis buffer, led to a 3.1-fold decrease in the fraction of mitochondrial reads compared to untreated ND samples (Fig. 3a, 0.16/0.05). However, unlike DT samples, the fraction of unique, non-mitochondrial reads increased only slightly (Fig. 3b, median fold-change: 1.1), probably because the fraction of mitochondrial reads was already small. Additionally, the effect of the anti-mt CRISPR treatment was inconsistent, with three samples showing a decrease in the fraction of usable reads and four showing an increase (Fig. 3b, dashed lines). When calling peaks in anti-mt CRISPR-treated ND samples, this inconsistency was also observed in some of the comparisons performed with different peak calling parametersand read depths, with some of the samples showing an increase in the number of peaks over their untreated counterparts, while other samples had the opposite effect (Supplemental Fig. S1).

When comparing the effect of the anti-mt CRISPR treatment between ND and DT samples, the former underperformed DT samples in terms of peaks called by 1.3-fold fewer peaks (median of 30,182 vs 40,682, respectively). Therefore, combining the anti-mt CRISPR treatment with removal of detergent from the lysis buffer did not provide substantial gains over the original protocol with detergent that was treated with anti-mt CRISPR.

In addition to the number of peaks called in the different treatments, we evaluated the quality of peaks using two other parameters: (i) the fraction of Gencode(8) transcription start sites (TSS’s) (Fig. 4a) and (ii) the fraction of Epigenome Roadmap(9) annotated enhancers overlapping peaks (Fig. 4b and Supplemental Fig. S2). The highest fraction of TSS’s and enhancers was identified in samples treated with anti-mt CRISPR, regardless of whether they were generated with or without detergent. Whereas both anti-mt CRISPR-treated DT and ND samples identified similar numbers of TSS’s, enhancers were identified at a higher rate using detergent in conjunction with the anti-mt CRISPR treatment. This difference could be explained by the notion that chromatin tends to be more open in promoters to allow transcription, while enhancers, due to their dynamic nature, would be less accessible. In this scenario, finding enhancers requires lower background and higher quality data, which we have shown is best represented by the detergent anti-mt CRISPR samples.

**Figure 4.**
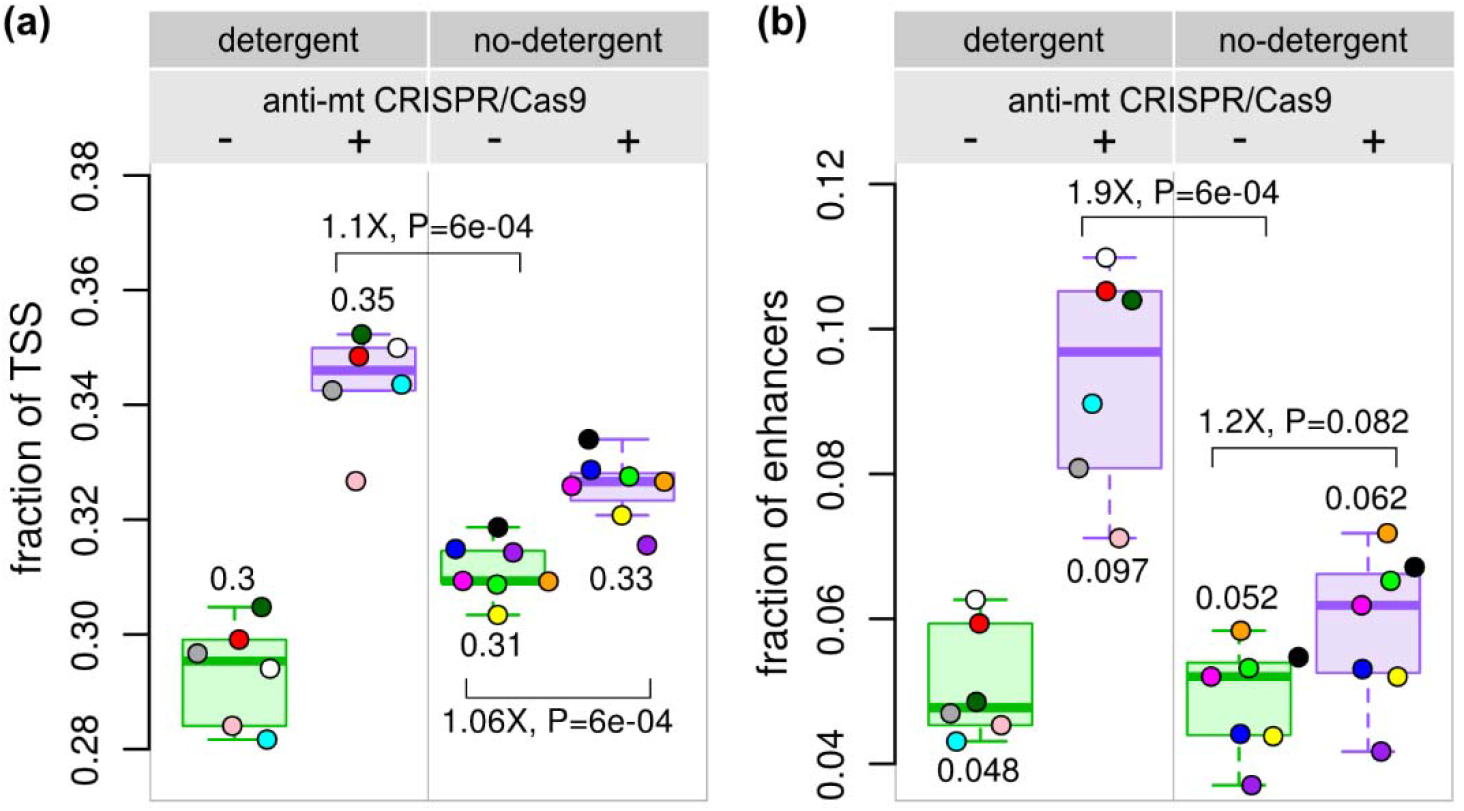
Comparison of the fraction of functional regions overlapping ATAC-seq peaks. (a) The fraction of transcription start sites (TSS’s) overlapping an ATAC-seq peak (+/−1 kb) was slightly higher in the DT samples than in the ND samples (1.05-fold). (b) Treated DT samples identified a greater number of Epigenome Roadmap GM12878 lymphoblastoid cell active enhancers (1.9-fold) than anti-mt CRISPR untreated ND samples. Fold-differences calculated on medians.

We conclude that reducing mitochondrial reads by cleavage of DNA sequencing fragments using an anti-mt CRISPR strategy yielded the best results in terms of numbers of peaks identified and their quality.

### Other treatments

Given the success of using CRISPR/Cas9 to reduce the amount of mitochondrial reads, we tested modifications of the treatment to enhance the degree of depletion of mitochondrial reads (Fig. 5). We tested (i) a longer Cas9 incubation of 2 hours instead of 1 hour, (ii) addition of Cas9 for an additional 1 hour after the initial 1 hour treatment (Cas9 boost) and (iii) adding 40X, 200X and 400X Cas9 instead of 100X. None of the treatments led to enhanced depletion of mitochondrial reads and, intriguingly, the 200X and 400X Cas9 treatments performed poorer than the 100X treatment (Fig. 5).

**Figure 5.**
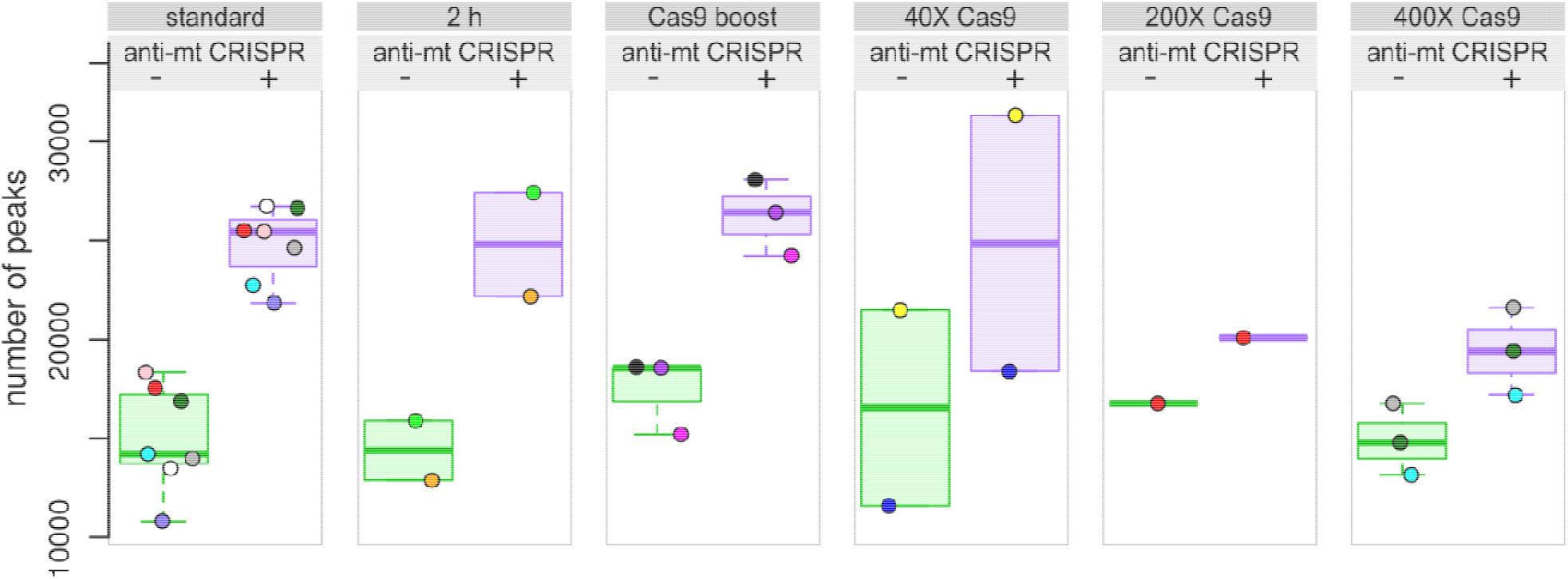
Modifications of the anti-mt CRISPR treatment. Compared to the treatment shown in Fig. 1 (100X gRNA, 100X Cas9, 1h incubation), labeled “standard”, modifications in the treatment did not show improvement. The number of peaks is comparable or even lower in the modified treatments, compared to the standard treatment. Due to the low number of reads in 6 samples, the results presented were obtained with 9.8 M reads randomly sampled. See also Supplemental Fig.S3.

We did not test a larger number of gRNAs targeting the mitochondrial genome, but it is likely that using 200 gRNAs instead of 100, for example, could further reduce the fraction of mitochondrial reads. However, as guide RNAs are priced per unit, the cost of the treatment increases linearly with the number of targets.

## Discussion

Our CRISPR/Cas9 treatment targeting 100 loci of the human mitochondrial chromosome successfully reduced the number of mitochondrial reads by 1.8-fold, similarly to Wu *et al*.(3), and increased the number of usable reads by 1.6-fold. Consequently, at the same read depth, samples generated with the original ATAC-seq protocol (DT) and treated with CRISPR/Cas9 and our anti-mt gRNAs resulted in 1.6-fold more peaks than the untreated controls. More TSS’s and enhancers were identified by peaks called in the treated samples, showing that the treatment increases the signal and does not induce unwanted changes in the data.

Removing detergent from the cell lysis step (ND) resulted in even lower number of mitochondrial reads (3.1-fold), but the peaks called were fewer and of lower quality. While the anti-mt CRISPR treatment improved ND samples, resulting in increased number of peaks called, it did not improve over DT samples. We observed more variability in treated ND samples than DT samples, as well as higher background, lower number of peaks and lower overlap with LCL enhancers.

In conclusion, our data show that treating samples prepared using detergent with gRNAs/Cas9 targeting mtDNA is the best way to reduce mtDNA contamination, increase the number of peaks, and improve identification of features such as TSS’s and enhancers.

Given the cost of gRNA oligos, sacrificing sequencing reads may be more economical than depleting mitochondrial reads if only a few samples are generated. gRNA oligos used in this work were purchased for US$ 2,300 in February, 2016, which was approximately the cost of three single-end 50 bp sequencing lanes on an Illumina HiSeq 4000 machine at the University of Chicago Functional Genomics Core Facility. The oligos can be amplified indefinitely and the Cas9 enzyme (US$ 540, New England Biolabs #M0386M) is enough for 25-30 samples, or about US$20 extra per sample. Laboratories could also split the cost and share the gRNAs to reduce the up-front investment in this treatment.

Caution should be taken when multiplexing anti-mt CRISPR-treated samples with samples that have not been treated. Treated samples will yield fewer sequencing reads unless a higher library concentration is used relative to other untreated samples. This is because the cleaved mitochondrial fragments will remain in the library but will not be sequenced since they cannot be amplified by bridge amplification. Sequencing a full lane of samples treated the same way does not require any adjustments.

During the execution of this project, an improved ATAC-seq method was published, termed Fast-ATAC(10), which uses a milder detergent in the cell lysis buffer. This treatment was reported to decrease the fraction of mitochondrial reads from 50% to 11%, while increasing the enrichment of reads in peaks over background and yielding more fragments per cell. The authors noted that cells that are more resistant to lysis may require a stronger detergent, i.e., the original ATAC-seq protocol, in which case, using the CRISPR treatment we analyzed here will remain useful. Since the cost of gRNAs is fixed and can be distributed among multiple laboratories, reducing mtDNA contamination using an anti-mt CRISPR treatment could still lead to significant savings if large numbers of samples are generated.

## Author Contributions

LEM proposed, planned and conducted experiments and edited the manuscript. LG designed the gRNAs. LG and ZZ helped perform experiments. YG contributed LCLs. GC and CO contributed ideas to the project. MAN conceived and supervised the project. NJS planned experiments, analyzed and interpreted the data and wrote the manuscript. All authors reviewed and approved the manuscript.

## Ethics Statement

The YRI cell lines were purchased from Coriell Cell Repository. The original samples were collected by the HapMap project in between 2001-2005. All of the samples were collected with extensive community engagement, including discussions with members of the donor communities about the ethical and social implications of human genetic variation research. Donors gave broad consent to future uses of the samples, including their use for extensive genotyping and sequencing, gene expression and proteomics studies, and all other types of genetic variation research, with the data publicly released.

